# Evaluation of endogenous miRNA reference genes across different zebrafish strains, developmental stages and kidney disease models

**DOI:** 10.1101/2021.04.09.439067

**Authors:** Florian Siegerist, Tim Lange, Anna Iervolino, Thor Magnus Koppe, Weibin Zhou, Giovambattista Capasso, Karlhans Endlich, Nicole Endlich

**Author notes:** Address for correspondence: Prof. Dr. rer. nat. Nicole Endlich, Friedrich-Loeffler-Str. 23c, 17487 Greifswald, Germany. The authors contributed equally to this work (Florian Siegerist, Tim Lange).

## Abstract

The majority of kidney diseases arise from the loss of podocytes and from morphological changes of their highly complex foot process architecture, which inevitably leads to a reduced kidney filtration and total loss of kidney function. It could have been shown that microRNAs (miRs) play a pivotal role in the pathogenesis of podocyte-associated kidney diseases. Due to their fully functioning pronephric kidney, larval zebrafish have become a popular vertebrate model, to study kidney diseases *in vivo*. Unfortunately, there is no consensus about a proper normalization strategy of RT-qPCR-based miRNA expression data in zebrafish. In this study we analyzed 9 preselected candidates dre-miR-92a-3p, dre-miR-206-3p, dre-miR-99-1, dre-miR-92b-3p, dre-miR-363-3p, dre-let-7e, dre-miR-454a, dre-miR-30c-5p, dre-miR-126a-5p for their capability as endogenous reference genes in zebrafish experiments.

Expression levels of potential candidates were measured in 3 different zebrafish strains, different developmental stages, and in different kidney disease models by RT-qPCR. Expression values were analyzed with NormFinder, BestKeeper, GeNorm, and DeltaCt and were tested for inter-group differences.

All candidates show an abundant expression throughout all samples and relatively high stability. The most stable candidate without significant inter-group differences was dre-miR-92b-3p making it a suitable endogenous reference gene for RT-qPCR-based miR expression zebrafish studies.

## Introduction

MicroRNAs (miRs) regulate protein expression by translational suppression via RNA interference. With their seed sequence they bind specifically to target mRNAs, thereby blocking translation and/or facilitating mRNA degradation^1^. In the recent past it could have been shown that microRNAs play a pivotal role in kidney development, physiology and pathology^2-4^.

Over the years, larval zebrafish have emerged a popular model organism, due to their small size, high reproductive potential and genetic accessibility^5^. The fact that zebrafish develop a fully filtrating glomerulus attached to a pair of tubules with high structural and molecular homology two days past fertilization (dpf) makes this model highly relevant for the investigation of human glomerular kidney diseases^6^. Most of these diseases arise from highly specialized glomerular cells, the podocytes. They are central players for glomerular permselectivity as they form the filtration slits together with neighboring podocytes. This is ensured by their complex interdigitating branching morphology. Interruptions in this morphology or the loss of podocytes subsequently leads to the development of nephrotic syndrome, a pathologic condition in which the size selectivity of the kidney filter gets impaired. Patients with nephrotic syndrome develop high-molecular weight proteinuria together with hypoproteinemic edema as typical characteristics^7^. A multitude of studies used larval zebrafish to model proteinuric glomerular diseases may it be with the help of morpholino-guided knockdown of podocyte genes^8^,^9^ or by treatment with podocyte-directed drugs^10-12^.

The relevance of miRs for glomerular pathophysiology has been proven several times. Studies have shown that the podocyte-specific knockout of the miR-processing ribonucleases Dicer and Drosha leads to proteinuria and glomerulopathy^13-15^. Beside their functional roles in the pathogenesis of all known types of glomerular diseases, miR deregulations can play an important role as biomarkers^16^,^17^.

Currently the gold standard in miR quantification is RT-qPCR^18^. For comprehensive analysis of RT-qPCR expression data, a reliable normalization strategy is required. The most widely applied method is the use of at least one endogenous reference gene^19^,^20^. The requirements for a suitable normalizer are sample type-independent abundant expression and a high stability throughout a certain sample set and it should not exhibit expressional differences between specific sub- and treatment groups.

Unfortunately, there is no consensus about a normalization strategy in zebrafish miR expression data. This is even more obvious when it comes to kidney research: Most studies make use of endogenous reference genes typically used in human and rodent studies. To the best of our knowledge there is no study specifically dealing with suitable endogenous normalizers focusing on zebrafish development and kidney disease models.

To address this, we analyzed the suitability of nine, from miR-sequencing preselected, miRs as endogenous reference genes for RT-qPCR based zebrafish miR expression data in different zebrafish strains, developmental stages and four different glomerular disease models.

## Materials and methods

### Zebrafish breeding

Zebrafish embryos were staged as described before^21^. Embryos were produced from timed matings as described^22^ and reared in E3 medium at 28.5°C in the dark with at least daily medium changes. Embryos of the following genotypes were used: AB/TÜ wildtype (ZDB-GENO-010924-10), Casper (mitfa^w2/w2^, mpv17^a9/a9^, ZDB-FISH-150901-6638) derived from double homozygous incrosses and Cherry (Tg(*nphs2*:GAL4); Tg(*UAS*:Eco.nfsB-mCherry) (mi1004Tg; rw0144Tg, ZDB-FISH-160601-2 ^23^)).

### Drug treatment

Embryos with a Tg(*nphs2*:GAL4), Tg(*UAS*:Eco.nfsB-mCherry) background obtained from double transgenic incrosses and selected for strong and homogenous mCherry fluorescence in podocytes at 72 hours past fertilization (hpf), were podocyte-depleted by treating embryos with 5 mM metronidazole (MTZ) dissolved in 0.1% DMSO in E3 medium or 0.1% DMSO in E3 as a vehicle control from 96 to 120 hpf as described before^11^. To induce focal and segmental glomerulosclerosis (FSGS)-like disease in zebrafish embryos, partial podocyte depletion was performed as described before^12^. Briefly, double transgenic Tg(*nphs2*:GAL4), Tg(*UAS*:Eco.nfsB-mCherry) embryos were treated with 80 µM MTZ in 0.1% DMSO in E3 for 48h starting at 96 hpf. For each treatment three independent biological replicates from individual clutches of embryos were set up.

### Morpholino Injections

Antisense morpholino oligonucleotides were produced by Gene-Tools (Philomath, OR, USA). Following beforehand established and published morpholino sequences were used (MO IDs http://www.zfin.org): MO3-wt1a (ZDB-MRPHLNO-071107-2), 5′-CACGAACATCAGAACCCATTTTGAG-3′^8^; MO1-nphs1 (ZDB-MRPHLNO-051102-1), 5’
s-CGCTGTCCATTACCTTTAGGCTCC-3‘^9^. Standard control morpholino targeting an intronic region of the human HBB gene: 5’-CCTCTTACCTCAGTTACAATTTATA-3’. Lyophilized morpholinos were reconstituted in ultrapure water to a stock concentration of 1 mM. Before injections, target and control morpholinos were diluted to 100 µM in injection solution containing 100 mM KCl, 10 mM HEPES pH 7.6 and 1% phenol red as a visual marker and incubated at 65°C for 5 min to dissolve precipitates. Per embryo, 2 nl were injected in the yolk of 1-2 cell-stage embryos. Injected embryos were collected in E3 and transferred to 10 cm petri dishes with fresh E3. Embryos were checked daily for viability and medium was changed twice a day. For each target, three independent clutches of embryos were injected.

### Sample generation

At the described endpoints, 20 embryos per group were collected in TRIreagent (Sigma Aldrich), homogenized using a tissue disruptor (MP-FastPrep-24, MPBiomedicals) with ceramic beads, snap-frozen in liquid nitrogen and stored at - 80°C upon RNA isolation. Total RNA isolation was performed as described per manufacturer’s description. The RNA pellet was eluted in DEPC treated ultrapure water and stored at -80°C upon further processing. RNA concentration and purity as 260/280 nm ratio was determined fluorometrically using an Eppendorf Biophotometer (Eppendorf, Hamburg, Germany).

### Candidate selection

Endogenous normalization candidates were selected from pretrials. Herein, smallRNA sequencing was performed from isolated glomeruli of zebrafish larvae (Tg(*nphs2*:GAL4); Tg(*UAS*:Eco.nfsb-mCherry), ZFIN: ZDB-FISH-160601-2 backcrossed to mitfa^w2/w2^) treated with MTZ and DMSO as described above^12^ with slight changes. Glomeruli were isolated at 6 dpf manually by micropipetting after slightly disrupting embryos in a tissue homogenizer. From the sequencing results we excluded miRNAs with less than 10 reads and p-values below 0.05 of pairwise comparisons between the treatment groups. The residual miRNAs were sorted by log_2_ fold changes between the two treatment groups. From these values we selected the most stable miRNAs as presented by values below 0.1 or above -0.1. This resulted in the following 9 candidates: dre-miR-92a-3p, dre-miR-206-3p, dre-miR-99-1, dre-miR-92b-3p, dre-miR-363-3p, dre-let-7e, dre-miR-454a, dre-miR-30c-5p, dre-miR-126a-5p.

### RT-PCR

For RT-PCR, 1 µg of RNA was reverse-transcribed to cDNA using the Quantitect Reverse Transcription Kit (Qiagen) according to manufacturer’s protocol. RT-PCR to assess MO knockdown efficiency was performed using the Phire Hot Start II DNA polymerase (Thermo Fisher Scientific) according to manufacturer’s instructions with 1 µl undiluted cDNA-template plus 19 µl master mix containing primers targeting exon 24-26 of *nphs1* and ef1a1l1 as a reference gene. Primer sequences were: nphs1_exon24_F: GTCTATGTGGTGGTGATCCTG, nphs1_exon26_R: CTGTGCCGAGGCGTTGATAA, ef1a1l1_F: AAGGAGGGTAATGCTAGCGG, ef1a1l1_R: GGGCGAAGGTCACAACCATA. As control a -RT, no template control from reverse-transcription and no template control from PCR Endpoint RT-PCR was run together with target samples in a Mastercycler gradient Thermocycler (Eppendorf) under following conditions: initial denaturation 98°C 3 min, 33 cycles: 98°C 10 s, 60°C 20 s, 72°C 25 s. RT-PCR products were resolved on 3.5% low melting agarose (Biozym) in 1x TBE-buffer containing 0.16 µg/ml ethidium bromide.

### Taqman™ miRNA Assay

Starting with 10 ng of total RNA, reverse transcription (RT) was performed using the Taqman™ miRNA Reverse Transcription Kit and Taqman™ miRNA Assays (Thermo Fisher Scientific). The following Taqman™ miRNA Assays were used: dre-miR-92a-3p Assay ID #000431; dre-miR-206-3p Assay ID #000501; dre-miR-99-1 Assay ID #000435; dre-miR-92b-3p Assay ID #007028_mat; dre-miR-363-3p Assay ID #001271; dre-let-7e Assay ID #005860_mat; dre-miR-454a Assay ID #007306_mat; dre-miR-30c-5p Assay ID #000419; dre-miR-126a-5p Assay ID #000451. The RT reaction was performed after manufacturer’s instruction using pooled primers. We included no template-as well as no reverse transcriptase controls. Additionally, a pooled RNA sample was used as positive control and inter-run calibrator that was synthesized with every RT run. RT-qPCR was performed using the above mentioned Taqman™ miRNA Assays and Taqman™ Universal Master Mix II, no UNG (Thermo Fisher Scientific) after manufacturer’s instructions. All samples were run in triplicate and each reaction consisted of 1.33 µL undiluted cDNA plus 18.7 µL master mix. An additional no template control was added to the negative controls from the RT reaction. The qPCR was performed on the Bio-Rad iCycler Thermal Cycler with the iQ5 Multicolor Real-Time PCR Detection System (Bio-Rad, Hercules, CA, USA) with the following cycler scheme: 10 min at 95°C initial denaturation; 45 cycles of 15 sec at 95°C and 60 sec at 60°C. The qPCR primary data analysis was done by the Bio-Rad iQ5 2.1 software with automatically set thresholds and baselines. Raw Ct-values ≥ 38 were excluded from analysis. All Ct-values were inter-run-calibrator corrected.

### Normalization analysis

The inter-run calibrator corrected values were analyzed for their stability by the online-based tool RefFinder^24^ (https://www.heartcure.com.au/reffinder/). It comprises the normalization tools BestKeeper^25^, comparative DeltaCt^26^, NormFinder^27^ and GeNorm^28^. These tools are based on different algorithms to evaluate the most stably expressed gene or gene pair of a specific sample set. For stability analysis we used different data sets: all values together, strains only, MTZ only and morpholinos only.

### Statistical analysis

Statistical analysis was performed with IBM SPSS 22.0 (SPSS Inc., Chicago, IL, USA) and GraphPad prism V5.01 (GraphPad Software, CA, USA). Data was checked for gaussian distribution by Kolmogorov-Smirnov test. All groups were tested for statistically significant differences by two-way ANOVA and Bonferroni *post-hoc* test. All values are displayed as means with standard deviations. P-values ≤ 0.05 were considered as statistically significant.

## Results

### Zebrafish kidney disease models: Morpholino-mediated gene knockdown

As established and widely used kidney disease models, we injected 1-cell-stage embryos with antisense morpholino oligonucleotides targeting *wt1a* translation initiation (translation blocking MOs: TBM) and *nphs1* splicing of the intron between *nphs1* exons 24-25 (splice blocking MOs: SBM). While the transcription factor *wt1a* plays an important role during glomerular development, *nphs1* is a main component of the mature glomerular filtration barrier. Generally, as shown in Fig.1A, injection of both MOs in 1-cell stage of zebrafish embryos resulted in typical edematous alterations frequently seen in glomerular damage due to high-molecular weight proteinuria and subsequent hypoproteinemia. Edema was graded in four categories from 0 – normal phenotype to 3 – severe whole-body edema with bent body axis. As shown in Fig.1B and C, a statistically significant proportion of embryos injected with 2 nl of 100 µM anti-*wt1a* or anti-*nphs1* morpholinos developed statistically significant pericardial or periocular edema. While the proportion of embryos showing edema of any severity was similar in both groups, the phenotype of the *nphs1* knockdown embryos was generally more severe compared to the *wt1a* knockdown (Fig. 1B). As shown in the exon-intron scheme in Fig.1D, the SBM used for the knockdown of *nphs1* blocks the splice donor site leading to a truncated protein due to integration of the intron between exons 24 and 25. RT-PCR (oligo positions depicted in Fig. 1D) amplifying exons 24-26 showed integration of the intron only after injection of the *nphs1* SBM (889 bp) with down-regulation of the wild type allele (210 bp) not containing the intron. As a reference gene *ef1a1l1* was used which was stably expressed across samples (Fig. 1E).

**Figure 1:**
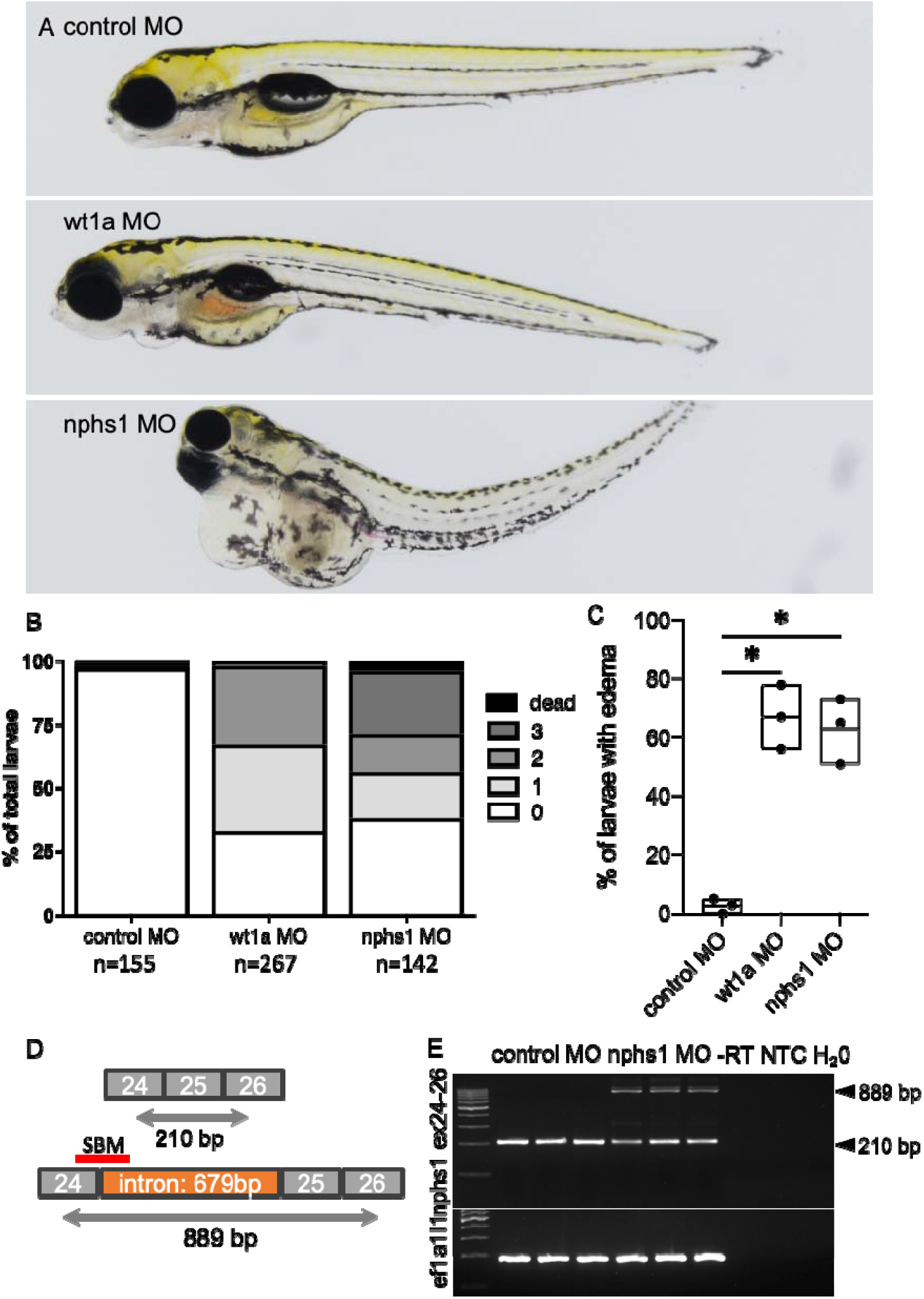
Induction of primary glomerular disease in larval zebrafish: Zebrafish embryos were injected with 2 nl 100 µM translation-blocking anti wt1a, splice-blocking anti-nphs1 or non-binding control MOs. A total of n=564 zebrafish embryos were injected and phenotyped at 120 hpf. Phenotype distribution graded in 0:no edema to 3: severe edema with bent body axis and dead larvae. As shown in the stacked boxplots in B, the phenotype of the wt1a knockdown larvae was generally more severe compared to nphs1 knockdown while the general proportion of larvae with edema was similar in both groups (C). No overt edema was seen in control MO injected larvae and the percentage of larvae with edema of any kind was statistically significant higher compared to paired control groups (paired t-test, p=0.01). D shows the exon-intron structure of exons 24-26 in the nphs1 gene with respective PCR-primer sites and the SBM MO binding site. Binding of the nphs1 SBM should lead to integration of a 679 bp intron. E shows agarose gel-resolved RT-PCR products for the nphs1 region described in D for three independent control and nphs1 SBM injected groups. Arrowheads show the size of the respective wildtype-bands at 210 and the shifted PCR product after intronic integration.

### Zebrafish kidney disease models: Pharmacogenetic podocyte depletion

Two protocols to injury adult podocytes were used: First, embryos expressing bacterial nitroreductase (NTR) specifically in podocytes were treated with a high concentration of MTZ (5 mM) from 4 to 5 dpf to deplete most podocytes from the GBM. This induces an acute onset of proteinuria mimicking acute nephrotic syndrome. As shown in the graph in Fig.2 A, 89% of larvae developed significant edema after 24 hours of treatment with 5 mM MTZ while only 4% of 0.1% DMSO control-treated larvae developed edema. This model leads to a rather acute form of podocyte injury and induces a rapid-onset nephrotic syndrome in zebrafish larvae.

In contrast to that, a lower concentration of MTZ depleted only a subset of podocytes and leads to a prolonged disease course. This mimics human FSGS with its diverse features (progressive proteinuria, parietal epithelial cell activation, extracellular matrix deposition) as we have shown before^12^. As shown in Figure 2 A, the phenotype progressively developed during the disease course over 4 days after the washout of MTZ with a lower concentration with increasing lethality until 9 dpf (Fig. 2B).

**Figure 2:**
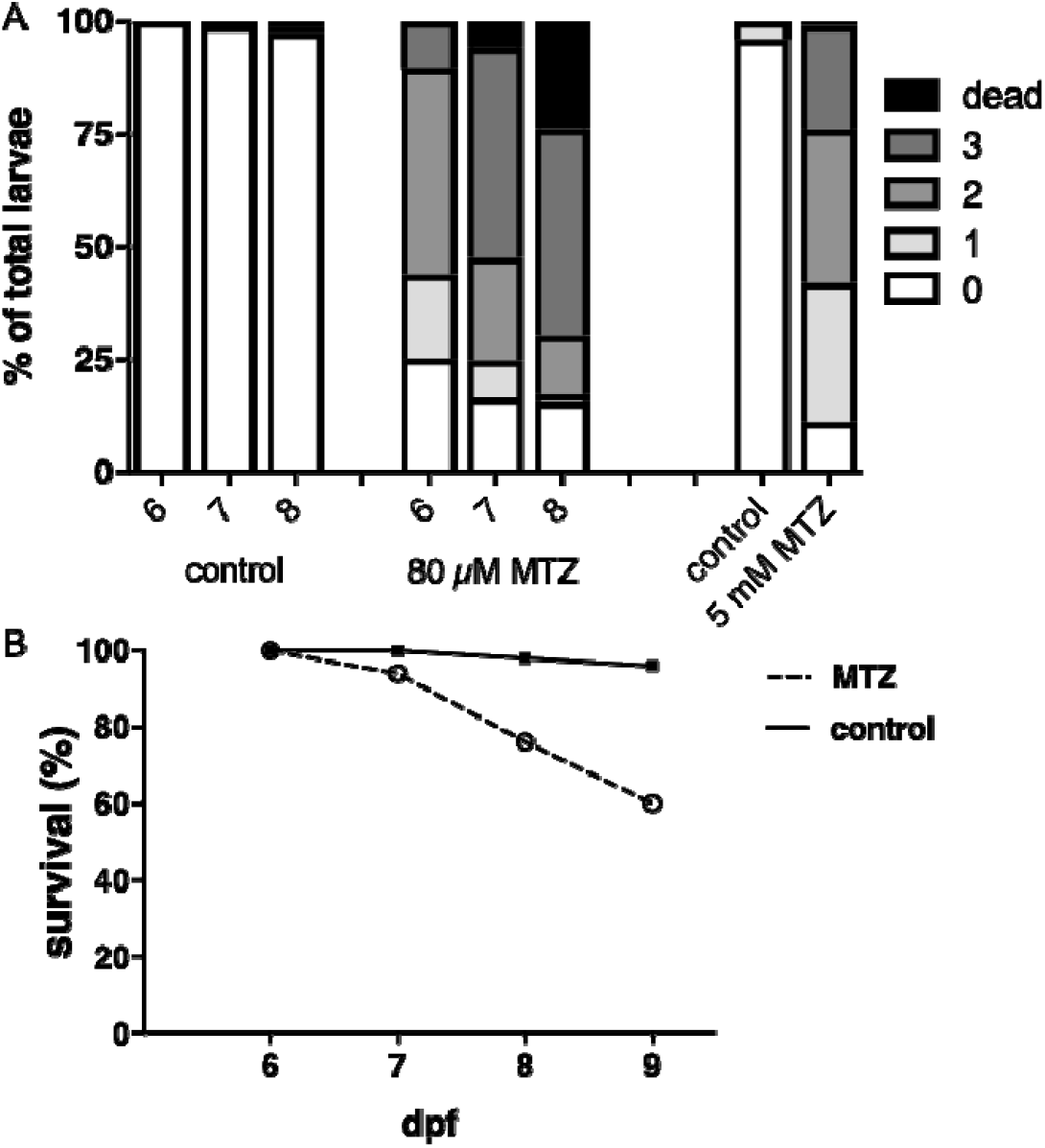
Acute nephrotic syndrome and focal and segmental Glomerulosclerosis zebrafish models: (A) Two models of podocyte depletion were used: First, 80 µM MTZ was applied over 48 hours. Edema developed in the proportions shown in A. As shown in B, mortality increased significantly 48 hours after MTZ washout.

## Expression of candidate as endogenous controls

We investigated the expression of dre-miR-92a-3p, dre-miR-206-3p, dre-miR-99-1, dre-miR-92b-3p, dre-miR-363-3p, dre-let-7e, dre-miR-454a, dre-miR-30c-5p and dre-miR-126a-5p by RT-qPCR. Mean Ct-values of the single samples were inter-run calibrator corrected. We observed an abundant expression of all candidate miRs in all samples. All candidate miRs showed a relatively homogenous expression pattern throughout the different strains and treatment groups (Fig 3). Dre-miR-92b-3p showed the lowest standard deviation (SD) of 0.018 followed by dre-miR-92a-3p (SD=0.23), dre-miR-206-3p (SD=0.029), dre-miR-363-3p (SD=0.030), dre-miR-99-1 (SD=0.032), dre-miR-30c-5p (SD=0.032), dre-miR-126a-5p (SD=0.035), dre-miR-454a (SD=0.036) and dre-miR-let7e (SD=0.047), respectively.

**Figure 3:**
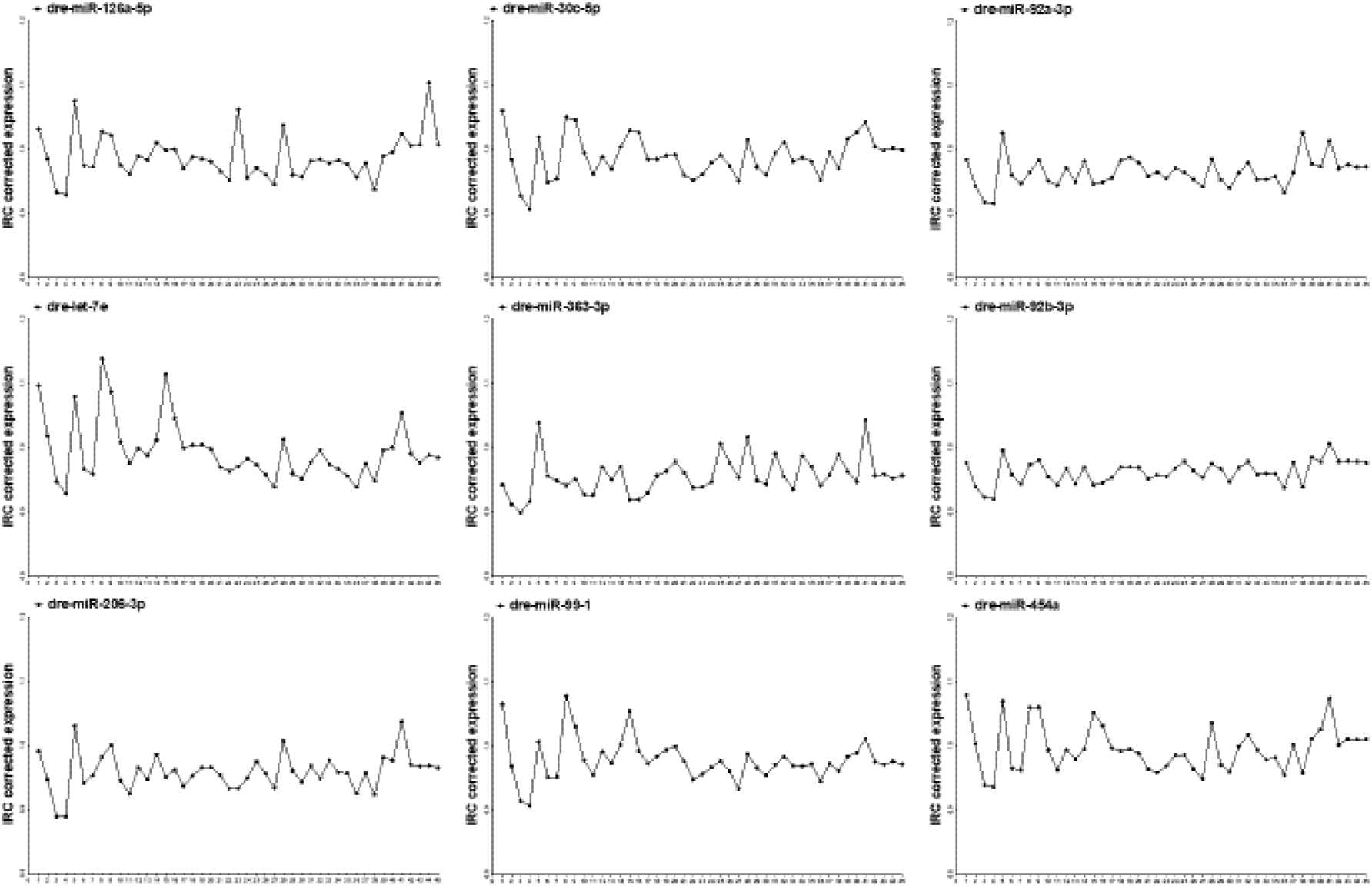
Abundant expression of candidate as endogenous control miRs. All miR candidates were detectable in the whole sample set. In general, they exhibit a high homogeneity as indicated by low standard deviations (SD). MiR-92b-3p shows most homogenous expression levels. Data is presented as inter-run calibrator (ICR) corrected expression. All values are shown as mean from technical triplicates. Numbers on the x-axis correspond to different strains, developmental stages and drug treatments.

### Stability analysis

The IRC corrected miRNA expression data was analyzed by the combined online tool RefFinder. For the analysis we used all unsorted values together as well as subclassifications such as only strains, only MTZ treatments and only morpholino-mediated knockdowns as input. The data was analyzed by BestKeeper, Genorm, Normfinder and DeltaCt.

#### BestKeeper

BestKeeper ranked the normalization candidates within the unsorted values from most stable to less stable as follows: dre-miR-92b-3p (0.015), dre-miR-92a-3p (0.017), dre-miR-206-3p (0.021), dre-miR-363-3p (0.021), dre-miR-99-1 (0.022), dre-miR-30c-5p (0.025), dre-miR-126a-5p (0.026), dre-miR-454a (0.028), dre-let-7e (0.034). dre-miR-126a-5p (0.023), dre-miR-30c-5p (0.03), dre-miR-99-1 (0.032), dre-miR-454a (0.033), dre-let-7e (0.044).

Looking at strains only, BestKeeper ranked the candidate miRs in the following order: dre-miR-92b-3p (0.016), dre-miR-92a-3p (0.020), dre-miR-363-3p (0.022), dre-miR-206-3p (0.023), dre-miR-126a-5p (0.023), dre-miR-30c-5p (0.030), dre-miR-99-1 (0.032), dre-miR-454a (0.033), dre-let-7e (0.044).

Based on the expression data obtained from MTZ treatments the ranking changed in the following way: dre-miR-92b-3p (0.009), dre-miR-92a-3p (0.011), dre-miR-99-1 (0.011), dre-let-7e (0.015), dre-miR-206-3p (0.016), dre-miR-30c-5p (0.016), dre-miR-454a (0.018), dre-miR-126a-5p (0.022), dre-miR-363-3p (0.023).

The RT-qPCR results from the morpholino-knockdown subset resulted in the following ranking: dre-miR-92b-3p (0.009), dre-miR-99-1 (0.010), dre-miR-30c-5p (0.015), dre-miR-92a-3p (0.016), dre-miR-454a (0.017), dre-let-7e (0.017),dre-miR-206-3p (0.018), dre-miR-363-3p (0.021), dre-miR-126a-5p (0.027) (Fig. 4).

**Figure 4:**
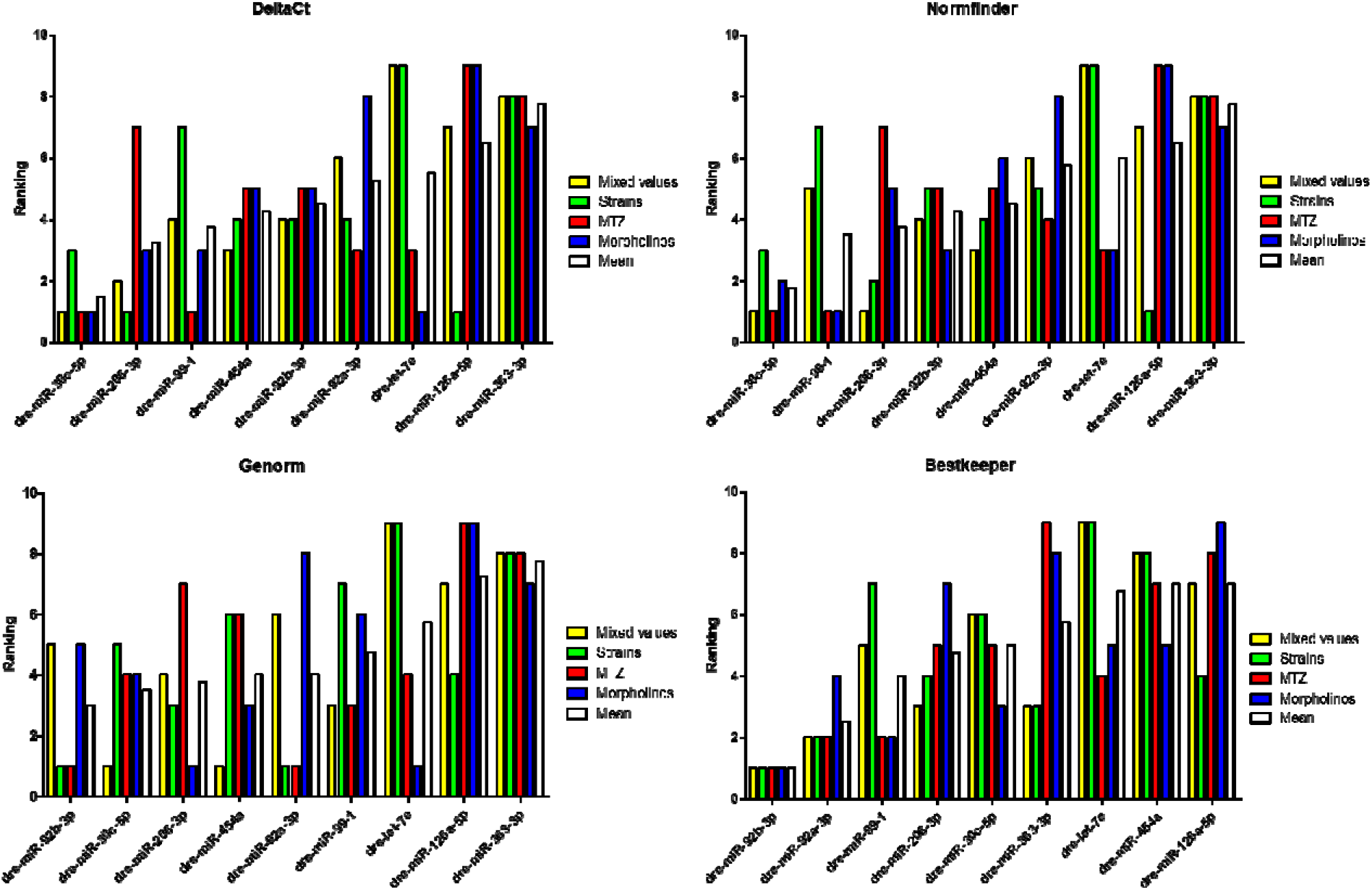
Ranking of potential endogenous control miRs by 4 different normalization determination softwares. Data was ranked by DeltaCt, Normfinder, Genorm and Bestkeeper with respect to mixed values, strains only, MTZ only and morpholinos only as well as the mean rank.

#### Genorm

The second normalization determination software Genorm ranked the miR candidates based on the mixed values as follows: dre-miR-454a (0.012), dre-miR-30c-5p (0.012), dre-miR-99-1 (0.015), dre-miR-206-3p (0.018), dre-miR-92b-3p (0.021), dre-miR-92a-3p (0.022), dre-miR-126a-5p (0.024), dre-miR-363-3p (0.026), dre-let-7e (0.028).

The stability ranking within the strains only showed this order: dre-miR-92a-3p (0.008), dre-miR-92b-3p (0.008), dre-miR-206-3p (0.014), dre-miR-126a-5p (0.015), dre-miR-30c-5p (0.02), dre-miR-454a (0.021), dre-miR-99-1 (0.023), dre-miR-363-3p (0.025), dre-let-7e (0.028).

Genorm ranked the miR candidates based on MTZ treatment expression values like this: dre-miR-92a-3p (0.007), dre-miR-92b-3p (0.007), dre-miR-99-1 (0.009), dre-let-7e (0.01), dre-miR-30c-5p (0.01), dre-miR-454a (0.011), dre-miR-206-3p (0.012), dre-miR-363-3p (0.015), dre-miR-126a-5p (0.018).

The potential endogenous controls within the morpholino subset were ranked as following: dre-miR-206-3p (0.005), dre-let-7e (0.005), dre-miR-454a (0.006), dre-miR-30c-5p (0.009), dre-miR-92b-3p (0.011), dre-miR-99-1 (0.012), dre-miR-363-3p (0.016), dre-miR-92a-3p (0.02), dre-miR-126a-5p (0.025) (Fig. 4).

#### Normfinder

With respect to the mixed values Normfinder software shows this ranking: dre-miR-206-3p (0.011), dre-miR-30c-5p (0.011), dre-miR-454a (0.014), dre-miR-92b-3p (0.016), dre-miR-99-1 (0.017), dre-miR-92a-3p (0.019), dre-miR-126a-5p (0.025), dre-miR-363-3p (0.03), dre-let-7e (0.032).

Normfinder ranked the potential normalizers with expression values from strains only as follows: dre-miR-126a-5p (0.007), dre-miR-206-3p (0.011), dre-miR-30c-5p (0.016), dre-miR-454a (0.017), dre-miR-92b-3p (0.019), dre-miR-92a-3p (0.019), dre-miR-99-1 (0.021), dre-miR-363-3p (0.032), dre-let-7e (0.035).

The MTZ-treatments resulted in the following order: dre-miR-99-1 (0.006), dre-miR-30c-5p (0.006), dre-let-7e (0.007), dre-miR-92a-3p (0.009), dre-miR-454a (0.01), dre-miR-92b-3p (0.01), dre-miR-206-3p (0.012), dre-miR-363-3p (0.02), dre-miR-126a-5p (0.029).

The ranking of the morpholino treatment group showed the following order: dre-miR-99-1 (0.005), dre-miR-30c-5p (0.007), dre-miR-92b-3p (0.008), dre-let-7e (0.008), dre-miR-206-3p (0.01), dre-miR-454a (0.012), dre-miR-363-3p (0.025), dre-miR-92a-3p (0.029), dre-miR-126a-5p (0.043) (Fig. 4).

#### DeltaCt

Delta Ct ranked the potential endogenous control miRs from mixed values as shown in the following: dre-miR-30c-5p (0.023), dre-miR-206-3p (0.024), dre-miR-454a (0.025), dre-miR-92b-3p (0.026), dre-miR-99-1 (0.026), dre-miR-92a-3p (0.027), dre-miR-126a-5p (0.032), dre-miR-363-3p (0.034), dre-let-7e (0.036).

When looking at the three different strains only the ranking was the following: dre-miR-126a-5p (0.023), dre-miR-206-3p (0.023), dre-miR-30c-5p (0.025), dre-miR-454a (0.026), dre-miR-92a-3p (0.026), dre-miR-92b-3p (0.026), dre-miR-99-1 (0.028), dre-miR-363-3p (0.035), dre-let-7e (0.038).

DeltaCt ranked the miR candidates based on MTZ-treatment values as follows: dre-miR-99-1 (0.014), dre-miR-30c-5p (0.014), dre-let-7e (0.015), dre-miR-92a-3p (0.015), dre-miR-92b-3p (0.016), dre-miR-454a (0.016), dre-miR-206-3p (0.018), dre-miR-363-3p (0.024), dre-miR-126a-5p (0.031).

The morpholino knockdown groups resulted in the following ranking: dre-let-7e (0.019), dre-miR-30c-5p (0.019), dre-miR-99-1 (0.02), dre-miR-206-3p (0.02), dre-miR-92b-3p (0.021), dre-miR-454a (0.021), dre-miR-363-3p (0.03), dre-miR-92a-3p (0.033), dre-miR-126a-5p (0.045) (Fig. 4).

#### Average ranking

For determination of the most stable normalization candidate miR, we calculated the mean rank for every single candidate within the specific determination software as well as the total mean. This revealed that dre-miR-30c-5p is the most stable candidate miR throughout the different treatment and sample types which got the lowest mean rank of 2.9. It was closely followed by dre-miR-92b-3p (3.2), dre-miR-206-3p (3.9), dre-miR-99-1 (4.0), dre-miR-92a-3p (4.4), dre-miR-454a (4.9), dre-let-7e (6.0), dre-miR-126a-5p (6.8) and dre-miR-363-3p as the least stable candidate miR with a mean rank of 7.3 (Fig. 5).

**Figure 5:**
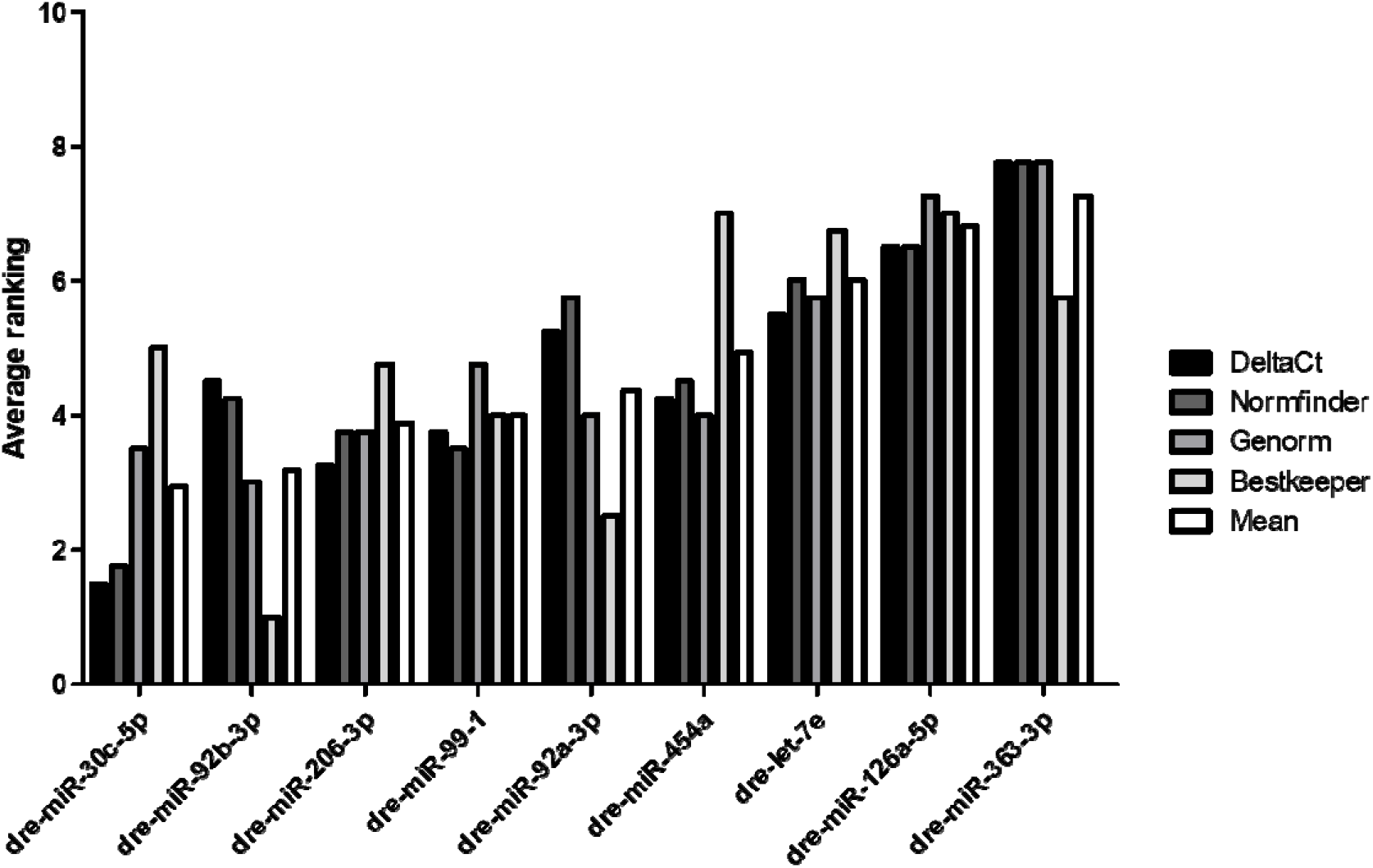
Average ranking of candidate miRs by 4 different normalization determination algorithms. Data was ranked by DeltaCt, Normfinder, Genorm and Bestkeeper.

### Inter-group differences

Since a certain endogenous control should not only be stable throughout a specific sample set but should not show significant expression differences between sample subgroups and treatment groups, we tested the expression data of the single miRs for the aforementioned differences. We could not find statistically significant expression differences between the three different strains for any tested miR. Additionally, there were no significant differences between the MTZ-treated groups themselves as well as between the treatment- and the control-groups. This was also true for the morpholino groups, where we could not detect any significant differences between the *nphs1*- and *wt1a* MOs groups as well as between them and the control group (Fig. 6).

**Figure 6:**
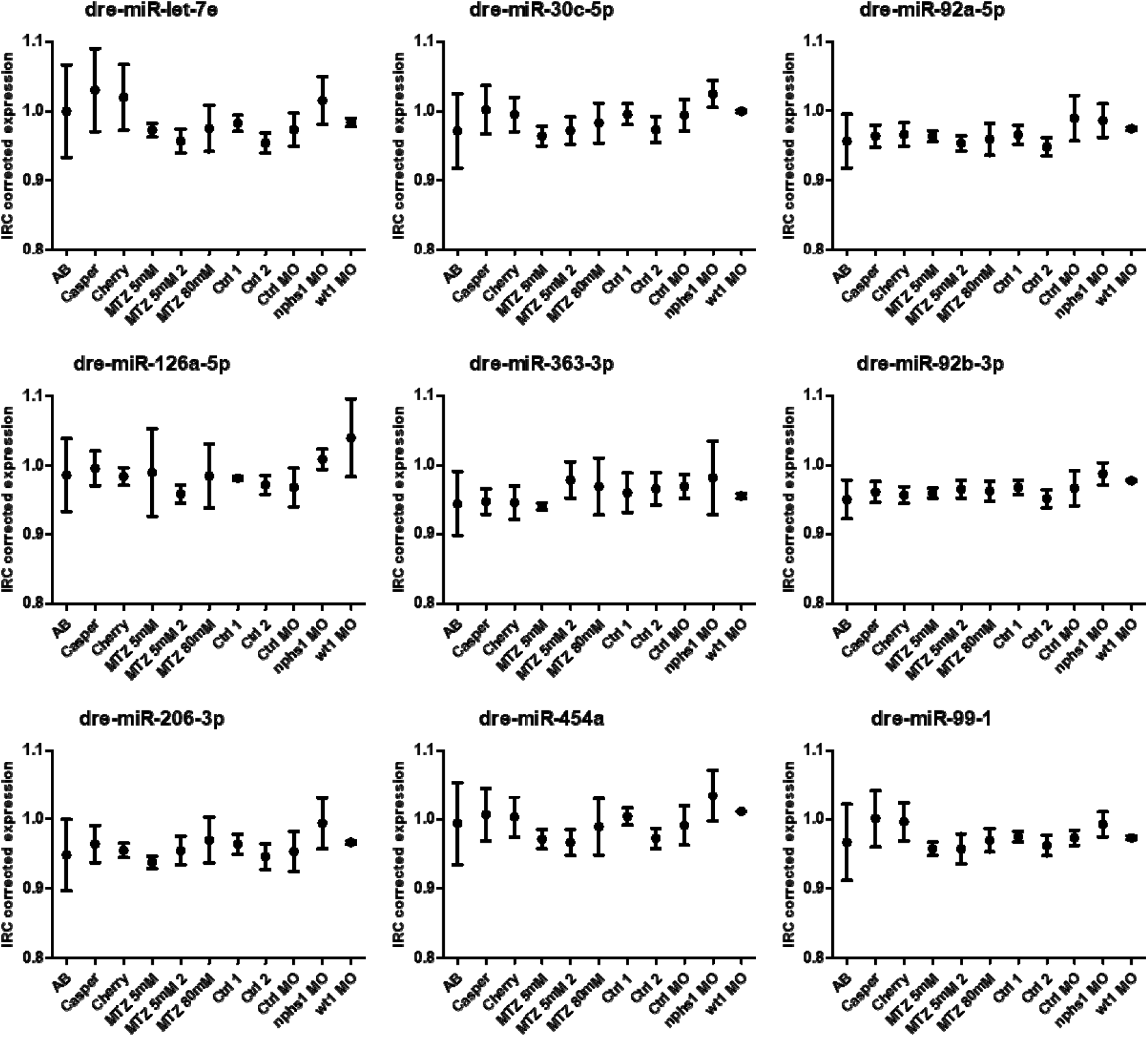
Differences in candidate miR expression between different subgroups. There are no significant differences between the presented subgroups. Data is presented as inter-run calibrator (ICR) corrected expression. All values are shown as mean from technical triplicates.

### Developmental differences

Beside specific developmental zebrafish research, many studies are dealing with zebrafish larvae at different developmental stages. In case of miR-based RT-qPCR data this fact sets the requirement for an endogenous control that shows either a high stability or no significant differences between developmental stages. To address this, we tested our data set for differences between the single developmental stages. For dre-miR-92a-3p, dre-miR-92b-3p and miR-126a-5p we could not detect any significant differences. The other six candidates showed differences between different developmental stages. Dre-miR-206-3p showed significant differences between 72 hours and 120 hours and between 96 hours and 120 hours. The next candidate, dre-miR-99-1, showed significant differences between 24 hours and the other time points. Dre-miR-363-3p had significant expression differences between 120 hours and 72 hours and between 120 hours and 48 hours. Dre-miR-let7e showed differences between 24 h and all other time points as well as between 48 h and 72 h, 96 h, 144 h and 192 h, respectively. The next candidate dre-miR-454a exhibited differences between 24 h and 72 h, 96 h, 144 h and 192 h, respectively, as well as between 48 h and 96 h. Interestingly, we could observe significant differences between developmental stages in the expression data of the most stably ranked normalization candidate dre-miR-30c-5p. It showed significant differences between 24 h and 72 h, 96 h, 144 h and 192 h, respectively, and between 48 h and 96 h (Fig. 7).

**Figure 7:**
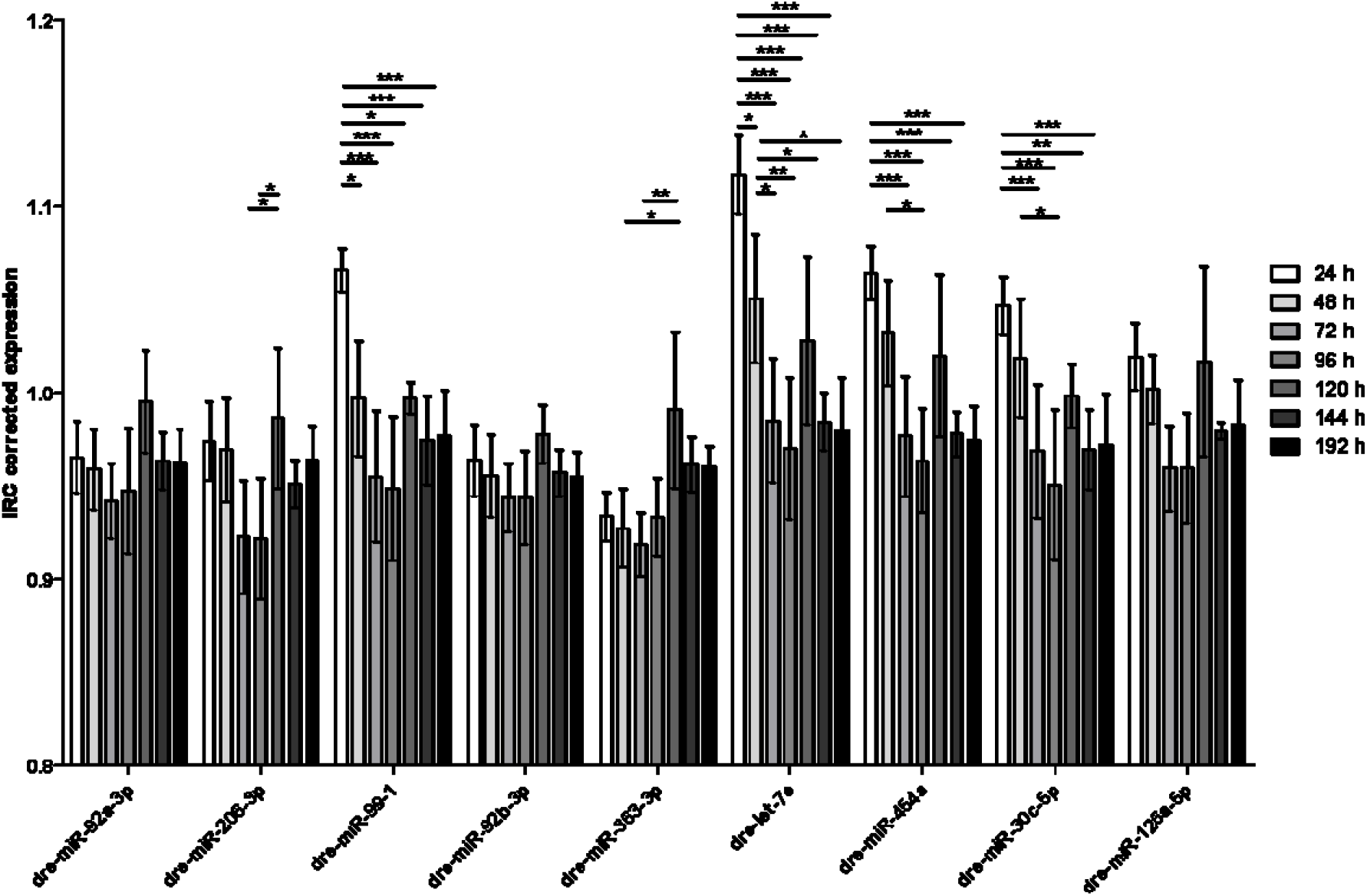
Differences in candidate miR expression between different developmental stages. There are no significant differences between the presented developmental stages in the expression of dre-miR-92a-3p, dre-miR-92b-3p and dre-miR-126a-5p. Data is presented as inter-run calibrator (ICR) corrected expression. All values are shown as mean from technical triplicates. * p ≤ 0.05, ** p ≤ 0.01, *** p ≤ 0.001.

## Discussion

Currently, there is no consensus about endogenous reference controls for RT-qPCR derived miR expression data in zebrafish larvae. To best of our knowledge, there are no studies explicitly approaching different drug treatments, knockdowns and/or developmental stages like it is the case for mRNAs^29^,^30^. In the present study we investigated the nine preselected candidates dre-miR-92a-3p, dre-miR-206-3p, dre-miR-99-1, dre-miR-92b-3p, dre-miR-363-3p, dre-let-7e, dre-miR-454a, dre-miR-30c-5p and dre-miR-126a-5p for their suitability as endogenous reference genes. As it has been shown before, zebrafish embryos exhibit highly dynamic miR expression patterns during development among different developmental stages in three different genetic zebrafish strains^2^,^31^. We started at 24 hours past fertilization (hpf), because lower time points have only a little relevance for kidney research^6^ and many miRs show no or only little expression below this time point or are influenced by the abundance of maternal miRs in the yolk^31^. Additionally, to different background strains and the developmental series, we applied four different models, successfully established to model glomerular diseases in zebrafish:

The first two models are based on the morpholino-guided knockdown of genes important for glomerular development and/or maintenance of the glomerular filtration barrier. The first one hindered translation initiation of *wt1a*, a transcription factor that regulates proper glomerular development and is still active in adult podocytes. The second one targeted proper splicing of *nphs1*, a gene encoding the protein nephrin which is a crucial part of the slit diaphragm and therefore maintaining glomerular filtration barrier function. In line with our results, knockdown of both genes has been shown to result in disrupted glomerular development as well as in early onset of high molecular weight proteinuria in larval zebrafish^8^,^9^.

The other models represent a pharmacogenetic form of highly specific podocyte depletion. They are grounded on the NTR / MTZ model of targeted tissue ablation which has been translated to podocyte research. In this model expression of the NTR, a bacterial enzyme, is transgenically driven by tissue-specific promotor fragments, such as in our case the *nphs2* promotor. When zebrafish embryos are treated with the antibiotic MTZ, the NTR converts this prodrug into a cytotoxin which leads to rapid apoptosis in the targeted tissue. Initially this model has been established to deplete pancreatic beta-cells and has later been modified to a dose-dependent depletion of podocytes^10^,^32^. We have shown that upon treatment with high dose MTZ (5 mM) zebrafish rapidly develop proteinuria and classic morphologic changes known from human nephrotic syndrome such as severe podocyte foot process effacement and later podocyte detachment^11^,^33^. Lately, we could show that when dose-dependently only a smaller subset of podocytes is depleted, a more chronic course of disease is initiated. In that model, zebrafish embryos resemble the phenotype known from human FSGS including activation of parietal epithelial cells that migrate to the glomerular tuft and deposit extracellular matrix^12^.

After RT-qPCR based measurements, the expression levels of candidate miRs showed a high homology in general in the described models. There were no statistically significant differences detectable between the single subgroups, except for developmental stages. The average ranking revealed dre-miR-30c-5p as the most stable endogenous reference gene. Dre-miR-30c-5p is known as a key player in pronephric development in *Xenopus laevis* and *Danio rerio*, where it shows a pronephros specific expression within the first days of development^34^,^35^. It has also been used as an endogenous reference gene before ^36^. Unfortunately, there were significant dre-miR-30c-5p expression differences between different developmental stages, meaning that it had to be rejected as a suitable normalizer for that kind of experimental setups. The only miRs that showed no differences in their expression levels between developmental stages were dre-miR-92a-3p, dre-miR-92b-3p and dre-miR-126a-5p. Dre-miR-92b-3p was the second most stable candidate closely after dre-miR-30c-5p. Additionally, it was exclusively ranked first in all subgroups by Bestkeeper. Dre-miR-92b-3p is known to be abundantly and ubiquitously expressed in zebrafish up from 24 hpf^37^ and is essential, together with dre-miR-92a, for the earliest steps in zebrafish embryogenesis ^38^.

The present study shows that all tested candidate miRs have a high homology within the tested experimental setups. The most stable normalizer was dre-miR-30c-5p. However, if it comes to developmental research studies or comparisons of zebrafish larvae at different ages it is rather unsuitable. Our analysis shows that dre-miR-92b-3p is the best candidate to be used as an endogenous reference gene for RT-qPCR-based miR expression in zebrafish larval studies.

